# A single-nucleus RNA sequencing atlas of the postnatal retina of the shark *Scyliorhinus canicula*

**DOI:** 10.1101/2024.04.12.589211

**Authors:** Nicolás Vidal-Vázquez, Ismael Hernández-Núñez, Pablo Carballo-Pacoret, Sarah Salisbury, Paula R. Villamayor, Francisca Hervas-Sotomayor, Xuefei Yuan, Francesco Lamanna, Céline Schneider, Julia Schmidt, Sylvie Mazan, Henrik Kaessmann, Fátima Adrio, Diego Robledo, Antón Barreiro-Iglesias, Eva Candal

**Affiliations:** Departamento de Bioloxía Funcional, Facultade de Bioloxía, Universidade de Santiago de Compostela, 15782, Santiago de Compostela, Spain; Aquatic One Health Research Center (ARCUS), Universidade de Santiago de Compostela, 15782, Santiago de Compostela, Spain; The Roslin Institute and Royal (Dick) School of Veterinary Studies, University of Edinburgh, Edinburgh EH25 9RG, UK; Departamento de Zooloxía, Xenética e Antropoloxía Física, Facultade de Veterinaria, Universidade de Santiago de Compostela, 27002, Lugo, Spain; Center for Molecular Biology (ZMBH), DKFZ-ZMBH Alliance, Heidelberg University, Heidelberg, Germany; INRAE, LPGP, Rennes, France; CNRS, Sorbonne Université, Biologie Intégrative des Organismes Marins, UMR7232-BIOM, Banyuls-sur-Mer, France; Departamento de Zooloxía, Xenética e Antropoloxía Física, Facultade de Bioloxía, Universidade de Santiago de Compostela, 15782, Santiago de Compostela, Spain

## Abstract

The retina, whose basic cellular structure is highly conserved across vertebrates, constitutes an accessible system for studying the central nervous system. In recent years, single-cell RNA-sequencing studies have uncovered cellular diversity in the retina of a variety of species, providing new insights on retinal evolution and development. However, similar data in cartilaginous fishes, the sister group to all other extant jawed vertebrates, are still lacking. Here, we present a single-nucleus RNA-sequencing atlas of the postnatal retina of the catshark *Scyliorhinus canicula*, consisting of the expression profiles for 17,438 individual cells from three female, juvenile catshark specimens. Unsupervised clustering revealed 22 distinct cell types comprising all major retinal cell classes, as well as retinal progenitor cells (whose presence reflects the persistence of proliferative activity in postnatal stages in sharks) and oligodendrocytes. Thus, our dataset serves as a foundation for further studies on the development and function of the catshark retina. Moreover, integration of our atlas with data from other species will allow for a better understanding of vertebrate retinal evolution.

## Background & Summary

The neural retina shows a remarkable degree of conservation in its cellular structure in all extant vertebrates^1–3^. This basic plan consists of five neuronal cell classes and one glial cell class, whose somata are arrayed in three nuclear layers, interspersed with two plexiform layers, where synapses occur (**Fig. 1a**). The outer nuclear layer (ONL) contains photoreceptors (PRs), the light-sensitive cells of the retina, which can usually be classified in two morphologically and functionally distinct major types, cones and rods, responsible for photopic and scotopic vision, respectively. In turn, the inner nuclear layer (INL) hosts three types of interneurons, namely horizontal cells (HCs), bipolar cells (BCs) and amacrine cells (ACs), which receive, integrate, modulate and transmit the signals coming from photoreceptors to retinal ganglion cells (RGCs). RGCs are located in the ganglion cell layer (GCL) and project their axons through the optic nerve to the visual processing centres of the brain. Besides, the retina contains a major glial cell class, Müller glia (MG), a type of radial glial cells whose nuclei are located within the INL. Other glial cell types, such as oligodendrocytes, microglia or astrocytes, may be present in the innermost layers of the retina, but, unlike MG, these are not derived from the optic cup; instead, they originate in other parts of the brain and migrate into the eye through the optic nerve^4^.

**Figure 1.**
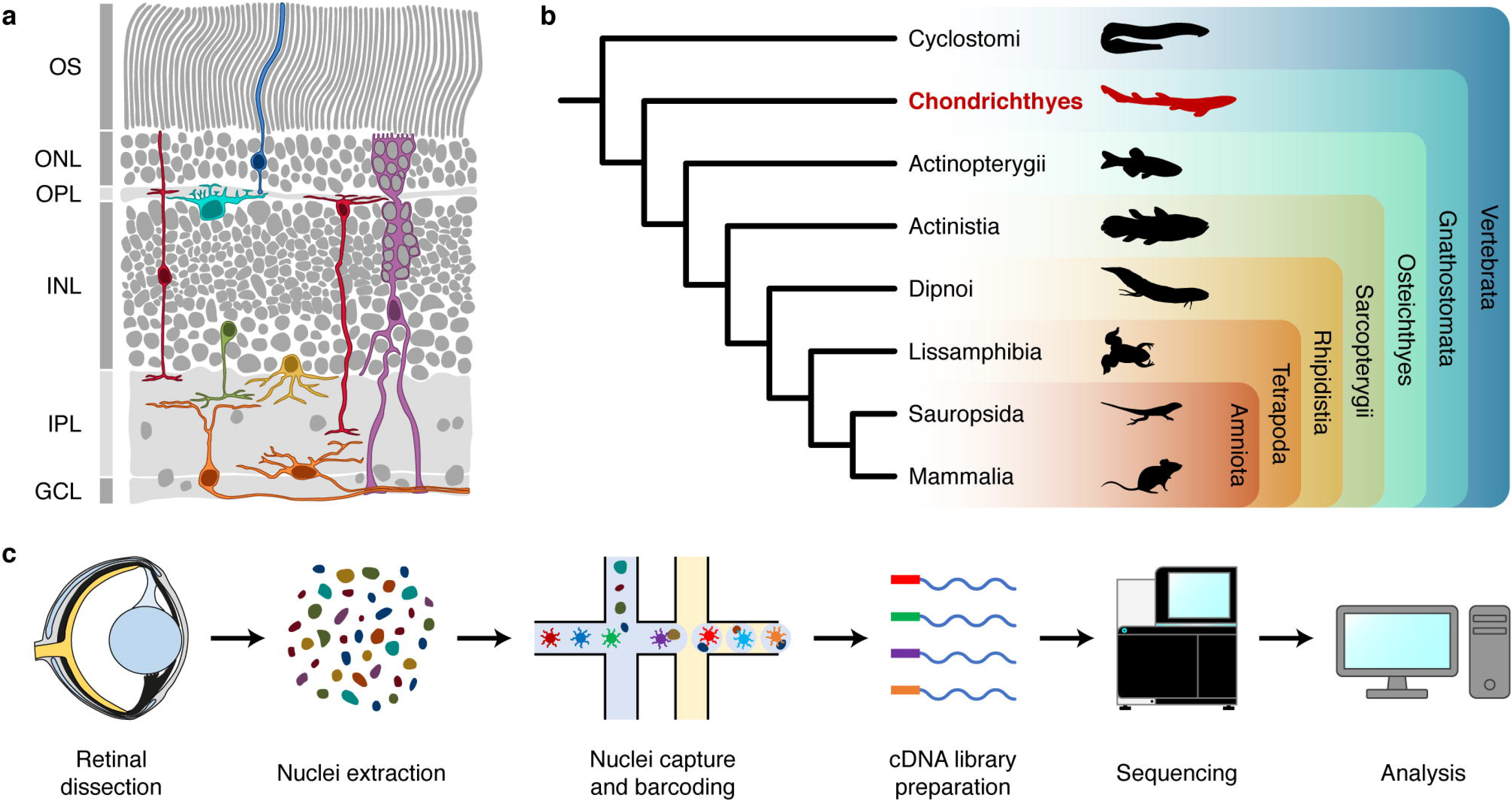
Background and experimental workflow. (**a**) Structure of the juvenile catshark retina, based on microscopy images kindly provided by Dr J. Francisco-Morcillo; cell drawings based on Neumayer^64^. (**b**) Cladogram showing relationships among major extant vertebrate groups, highlighting the position of chondrichthyans as the sister group to all other gnathostomes. Silhouettes from PhyloPic^84^. (**c**) Experimental workflow. Eye diagram after Collin^42^. GCL, ganglion cell layer; INL, inner nuclear layer; IPL, inner plexiform layer; ONL, outer nuclear layer; OPL, outer plexiform layer; OS, photoreceptor outer segments.

Most of these cell classes can be subdivided into a number of cell types with diverse morphological, functional and molecular properties^5–7^. In recent years, high-throughput single-cell RNA sequencing (scRNA-seq) studies have provided unprecedented insights into molecular cell type diversity in the mature and developing vertebrate retina^6,8–10^. Although most studies have focused on the retina of mammals^11–32^ (particularly that of rodents^12–18^ and primates^19–32^), cellular atlases of the chick^13,33^, brown anole lizard^11^, zebrafish^9,13,34–40^ and sea lamprey^11,41^ retinas have also been generated to date, providing new resources to study the evolution and development of vertebrate retinal cell types. However, scRNA-seq studies on the retina of cartilaginous fishes are still lacking.

Given their phylogenetic position as the sister group to all other extant gnathostomes (**Fig. 1b**), chondrichthyans (sharks, rays and chimaeras) constitute a particularly interesting group to study retinal evolution^42–44^. Among cartilaginous fishes, the shark *Scyliorhinus canicula* (Linnaeus, 1758), also known as the small-spotted catshark or the lesser-spotted dogfish, stands as a suitable model for experimental studies, owing to its abundance, relatively small size and accessible oviparous development^43^. Furthermore, the catshark retina shows persistent cell proliferation in postnatal stages^45,46^ (similar to other fishes^47^), making *S. canicula* an interesting species for the study of postnatal retinal neurogenesis.

In this study, we generated a single-nucleus RNA-sequencing (snRNA-seq) atlas of the postnatal retina of *S. canicula* (**Fig. 1c**). Early juvenile specimens were chosen to allow profiling of both mature and progenitor cell types, since previous work has shown that cell proliferation decreases as the animal grows, with mitotic activity being virtually absent in sexually mature adults^45,48^. Thus, we used three retinas from three female juvenile catsharks to generate a dataset consisting of 17,438 high quality nuclei. Unsupervised clustering revealed 22 cell types representing all major retinal cell classes, as well as retinal progenitor cells and oligodendrocytes. This constitutes, to the best of our knowledge, the first single-cell transcriptomic study of the retina of a chondrichthyan, providing a groundwork for comparative studies on the evolution of both retinal cell type diversity and retinal neurogenesis.

## Methods

### Animals

Three female, juvenile specimens of *S. canicula*, with a total length of 10.5 to 11.1 cm (**Table 1**) were kindly provided by the Aquarium Finisterrae in A Coruña (Spain) and kept in artificial seawater tanks under standard conditions of temperature (15-16 °C), pH (7.5-8.5) and salinity (35 g/L). All procedures were performed in accordance with the guidelines for animal experimentation established by the European Union and the Spanish government and were approved by the Bioethics Committee of the University of Santiago de Compostela (license number 15004/2022/001).

**Table 1.** Sample information.

| Sample | Total length (cm) | Sex |
| --- | --- | --- |
| Sample 1 | 11.0 | Female |
| Sample 2 | 10.5 | Female |
| Sample 3 | 11.1 | Female |

### Retina sampling

Animals were deeply anaesthetised with 0.5% tricaine methanesulfonate (MS-222; Thermo Fisher Scientific, 118000500) in seawater. The animals were removed from water, the eyes enucleated, and the retinas (one from each specimen, n = 3) were dissected out under a stereomicroscope. Retinas were immediately put in Eppendorf tubes and frozen in liquid nitrogen. Retinal samples were kept at −80 °C until they were further processed for nuclei isolation and RNA sequencing.

### Nuclei isolation and snRNA-seq data generation

Retina nuclei were extracted following a published protocol^49^ with small modifications. The frozen retinas were homogenised using a micropestle in 400 μL ice-cold homogenisation buffer (250 mM sucrose, 25 mM KCl, 5 mM MgCl_2_, 10 mM Tris-HCl [pH 8], 0.1% IGEPAL, 1 μM dithiothreitol [DTT], 0.4 U/μL Murine RNase Inhibitor [New England BioLabs, M0314L], and 0.2 U/μL SUPERase-In [Ambion, AM2694]). The homogenates were triturated gently using a p1000 tip for 10 times, incubated on ice for 5 min and then centrifuged at 100 g for 1 min at 4 °C to pellet any unlysed tissue chunks. The supernatant was transferred into another 1.5 mL Eppendorf tube and centrifuged at 400 g for 4 min at 4 °C to collect nuclei. The nuclei were washed twice in 400 μL homogenisation buffer and strained using a 40 μm Flowmi strainer (Sigma, BAH136800040) during the second wash step to remove nuclei aggregates. The final nuclei pellet was resuspended in 30-50 μL Nuclei Buffer (10x Genomics, PN-2000207). To estimate the nuclei concentration, nuclei aliquots were diluted in phosphate-buffered-saline (PBS) with Hoechst and propidium iodide (PI) DNA dyes and counted on Countless II FL Automated Cell Counter (Thermo Fisher Scientific, RRID: SCR_020236). Around 15,000 nuclei were used as input for the snRNA-seq experiment. The Chromium Next GEM Single Cell 3’ Reagent Kits v3.1 (PN-1000121, PN-1000120, and PN-1000213) were used to make snRNA-seq libraries. Libraries were quantified on a Qubit Fluorometer (Thermo Fisher Scientific; RRID: SCR_018095) and quality checked on a Fragment Analyzer (Agilent; RRID: SCR_019417). Libraries were sequenced on NextSeq550 (Illumina; RRID: SCR_016381; 28 cycles for Read 1, 56 cycles for Read 2, 8 cycles for i7 index).

### Genome indexing and read alignment

Genome indexing and library mapping was performed with STAR (v2.7.10a)^50^. The *S. canicula* genome assembly sScyCan 1.1^51^ (GCF_902713615.1; *GCF_902713615*.*1_sScyCan1*.*1_genomic*.*fna*) and its associated annotation in GFF format (*GCF_902713615*.*1_sScyCan1*.*1_genomic*.*gff*) were downloaded from the National Center for Biotechnology Information (NCBI). The mitochondria (NC_001950.1) annotations within the GFF file were manually edited to convert “CDS” annotations to “exon” annotations and to convert all annotations of “tRNA” and “rRNA” to “gene” annotation. This was done to ensure mitochondrial gene annotations were consistent with those of the nuclear genes as STAR assigns transcripts to “exon” annotations in the .gff file. The GFF annotation file was converted to GTF format using gffread (v0.10.1)^52^. The genome and its annotation (GTF) were indexed using STAR (--runMode genomeGenerate). Each library was then mapped against the genome with the 10x V3 cell barcode whitelist (*3M-february-2018*.*txt*) and using standard parameters for single cell libraries (--soloMultiMappers Unique --soloBarcodeReadLength 28 --soloType CB_UMI_Simple --soloUMIlen 12 -- soloCBwhitelist 3M-february-2018.txt --soloFeatures GeneFull --clipAdapterType CellRanger4 --outFilterScoreMin 20 --soloCBmatchWLtype 1MM_multi_Nbase_pseudocounts --soloUMIfiltering MultiGeneUMI_CR --soloUMIdedup 1MM_CR --readFilesCommand zcat --outSAMtype BAM Unsorted). The raw (unfiltered) files (*genes*.*tsv, barcodes*.*tsv*, and *matrix*.*mtx*) generated for each sample were then used for downstream analysis. On average, there were 274 million reads per sample with 94% of reads with valid barcodes, and a 45% saturation. A summary of the STAR output statistics for each sample can be found in **Table 2**.

**Table 2.** Summary statistics for STAR outputs for each sample. Note that raw files were used for downstream analysis (which includes all barcodes, not just those estimated to be viable cells by STAR). Therefore, the statistics relating to STAR-assigned cells are not necessarily relevant but are provided for the reader’s interest.

| Sample | Sample 1 | Sample 2 | Sample 3 |
| --- | --- | --- | --- |
| Number of reads | 229,861,905 | 249,344,842 | 343,563,083 |
| Reads with valid barcodes | 0.960049 | 0.932741 | 0.917902 |
| Sequencing saturation | 0.502799 | 0.521001 | 0.338812 |
| Q30 bases in CB + UMI | 0.957978 | 0.95938 | 0.961753 |
| Q30 bases in RNA read | 0.89873 | 0.90096 | 0.888598 |
| Reads mapped to genome: unique + multiple | 0.88851 | 0.846251 | 0.647372 |
| Reads mapped to genome: unique | 0.781012 | 0.711432 | 0.518868 |
| Reads mapped to GeneFull: unique + multiple<br>GeneFull | 0.693977 | 0.471452 | 0.354986 |
| Reads mapped to GeneFull: unique GeneFull | 0.642748 | 0.424422 | 0.314171 |
| Estimated number of cells | 3,028 | 1,074 | 2,521 |
| Unique reads in cells mapped to GeneFull | 62,372,962 | 34,713,678 | 39,214,416 |
| Fraction of unique reads in cells | 0.422171 | 0.328021 | 0.363307 |
| Mean reads per cell | 20,598 | 32,321 | 15,555 |
| Median reads per cell | 15,254 | 21,413 | 11,331 |
| UMIs in cells | 30,760,018 | 16,433,368 | 25,334,077 |
| Mean UMIs per cell | 10,158 | 15,301 | 10,049 |
| Median UMIs per cell | 7,593 | 10,144 | 7,340 |
| Mean GeneFull per cell | 3,508 | 4,872 | 3,988 |
| Median GeneFull per cell | 3,170 | 4,285 | 3,540 |
| Total GeneFull Detected | 23,271 | 23,076 | 23,449 |

### Bioinformatic quality control

Samples were then analysed in an R (v4) environment using Seurat (v4.3.0)^53^. We created Seurat objects for each library after removing nuclei with less than 200 features and features occurring in fewer than three nuclei. Nuclei where mitochondrial (mtDNA) features accounted for 10% or more of their total unique molecular identifiers (UMIs) were removed before removing all mtDNA features. After sub-setting the Seurat object into individual samples, upper and lower thresholds for UMI and feature counts per nuclei were then applied individually to each sample based on knee plot visualisation. In sample 1, nuclei with more than 1,000 but less than 20,000 UMIs and more than 500 but less than 5,500 features were retained. In sample 2, nuclei with more than 500 but less than 24,000 UMIs and more than 500 but less than 6,500 features were retained. Finally, in sample 3, nuclei with more than 750 but less than 20,000 UMIs and more than 300 but less than 7,000 features were retained.

Samples were then merged into a single Seurat object before splitting samples again into individual sample datasets. This was done to ensure that the same features were considered across samples. Counts were then normalised for each sample using the “NormalizeData” function prior to calculating cell cycle scores using the “CellCycleScoring” function (see **Supplementary Table 1** for list of genes used). The “v2” SCTransform version with the glmGamPoi method (v1.9.0)^54^ was used to normalise RNA counts for each sample, regressing out scores for the S and G2M cell cycle stages. Linear dimension reduction was conducted for each sample using the “RunPCA” function with 50 PCs. After consulting Elbowplots for each sample, a Uniform Manifold Approximation and Projection (UMAP) using 20 principal components (PCs) was run for each sample and the “FindNeighbours” function was applied using 20 PCs, before using the “FindClusters” function with a resolution of 0.5. DoubletFinder (v2.0.3)^55^ was then applied independently to each sample selecting pK values with the highest associated mean-variance normalised bimodality coefficient (BCmvn) value. We assumed a 4% doublet formation rate (based on the Chromium instrument specifications) and adjusted for homotypic doublets.

### Clustering and differential gene expression analyses

Samples were integrated using 2000 features and anchors that were identified with the “rpca” reduction method and the “FindIntegrationAnchors” function. A principal component analysis (PCA) was rerun on the integrated dataset using 50 PCs, and 30 PCs were used for subsequent UMAP generation and clustering with a resolution of 0.5. Markers for each cluster were assessed using the logistic regression method and the FindAllMarkers function on the “SCT” assay and “data” slot, using sample ID as a latent variable to help reduce batch effects among samples. We used a pseudocount of 0.001, set a p-value threshold of 0.01, and only considered genes that were upregulated, expressed in at least 25% of all nuclei (in either of the compared groups), and demonstrated a threshold of 0.25 X difference (log-scale) between the two compared groups. Three clusters that were mostly composed of cells from a single sample and did not show differential expression of typical retinal cell marker genes were removed (see below; **Supplementary Table 2** shows differentially expressed genes in the removed clusters), and all remaining cells were re-clustered using the same parameters. The same differential expression analysis was applied to various groups of clusters (**Table 3**) to identify marker genes for cell classes composed of several clusters.

**Table 3.**
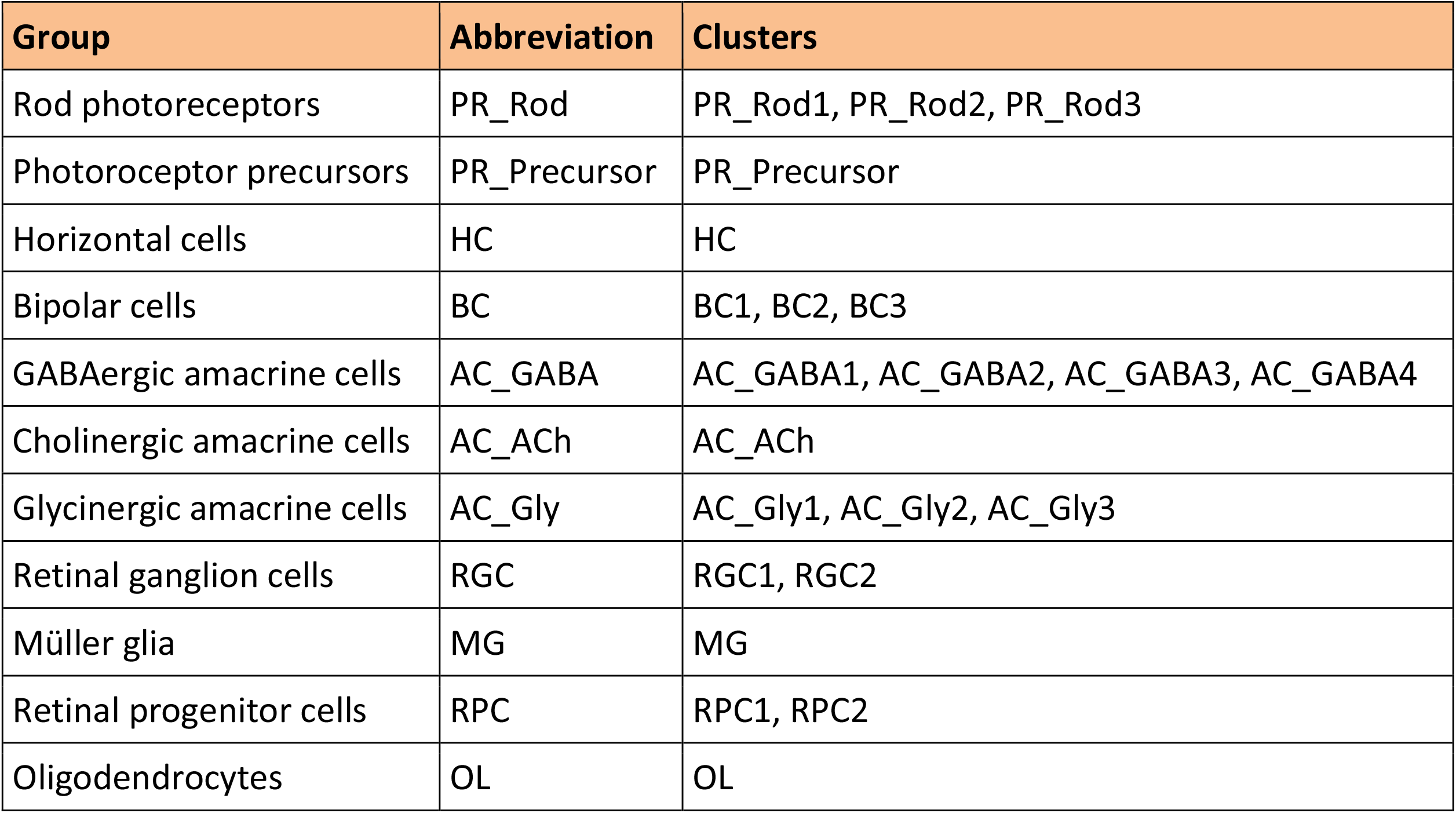
Class/subclass-level cluster groups for differential expression analysis.

| Group | Abbreviation | Clusters |
| --- | --- | --- |
| Rod photoreceptors | PR_Rod | PR_Rod1, PR_Rod2, PR_Rod3 |
| Photoreceptor precursors | PR_Precursor | PR_Precursor |
| Horizontal cells | HC | HC |
| Bipolar cells | BC | BC1, BC2, BC3 |
| GABAergic amacrine cells | AC_GABA | AC_GABA1, AC_GABA2, AC_GABA3, AC_GABA4 |
| Cholinergic amacrine cells | AC_ACh | AC_ACh |
| Glycinergic amacrine cells | AC_Gly | AC_Gly1, AC_Gly2, AC_Gly3 |
| Retinal ganglion cells | RGC | RGC1, RGC2 |
| Müller glia | MG | MG |
| Retinal progenitor cells | RPC | RPC1, RPC2 |
| Oligodendrocytes | OL | OL |

### Cluster annotation

To assign a cell class identity to each cluster, we elaborated a list of marker genes reported in previous studies on other vertebrate species^11–15,18–21,23,25–31,33,35,41^. Putative orthologues in the *S. canicula* genome were identified by comparing the sequence of the mouse gene (genome assembly GRCm39, GCF_000001635.27)^56^ with the *S. canicula* genome assembly sScyCan1.1 (GCF_902713615.1)^51^ using Blastn^57^. For marker genes absent from the mouse genome, either the human (genome assembly GRCh38.p14, GCF_000001405.40)^58^ or zebrafish (genome assembly GRCz11, GCF_000002035.6)^59^ genes were used. All marker gene sequences were obtained from GenBank^60^. Marker genes present in the *S. canicula* genome where then visualised in our dataset using the “FeaturePlot” and “DotPlot” functions of Seurat. This, coupled with the examination of the top differentially expressed genes in each cluster (see above), allowed us to assign all clusters in the final object to known retinal cell classes (see Technical Validation).

### Gene nomenclature

For the sake of readability, we decided to change the symbols of genes annotated with a LOC number in the main text and figures of the present article. We replaced LOC numbers for symbols based on the identification of their protein products, adding an “-*l*” at the end for those annotated as “-like”, except for uncharacterised genes, which were left with their LOC number. **Supplementary Table 3** shows the correspondence between the symbols used in the text and the sScyCan1.1 annotation, which is the one used in the dataset.

It should also be noted that many *S. canicula* genes follow the nomenclature used in zebrafish, including a letter or number at the end of their symbol. Since teleosts experienced an additional whole genome duplication after diverging from cartilaginous fishes^61^, many gene paralogues present in zebrafish are absent from the catshark genome. Therefore, current gene symbols do not accurately reflect the existence of paralogues in *S. canicula*, nor their correspondence to specific zebrafish paralogues.

## Data Records

Raw sequencing data for each sample (.*fastq* files) have been uploaded to the NCBI Sequence Read Archive (SRA) under BioProject accession number PRJNA1056918^62^. For each sample, STAR raw output files, i.e. expression matrices for each gene in each cell (*matrix*.*mtx*), barcodes (*barcodes*.*tsv*) and genes (*features*.*tsv*), have been uploaded to Figshare (doi: XXX). The final annotated Seurat object (*Scyliorhinus_retina*.*rds*) is also available on Figshare.

Two .*xlsx* files containing the list of differentially expressed genes in each cluster and each cell class (see **Table 3**) have been uploaded to Figshare. For each gene, the p-value (“p_val”), adjusted p-value (“p_val_adj”), average log_2_ fold change (“avg_log2FC”), the percentage of cells expressing that gene in the present cluster (“pct.1”) and the percentage of cells expressing said gene in the rest of the dataset (“pct.2”) are provided.

## Technical Validation

### Quality control

Retinas from three similarly sized, female, juvenile *Scyliorhinus canicula* specimens (**Table 1**) were used to generate the snRNA-seq dataset. To ensure that all the obtained barcodes correspond to viable nuclei, we established selection criteria based on feature counts, UMI counts and expression of mitochondrial genes (**Fig. 2a, b**). Prior to filtering, the three samples showed a similar number of nuclei (**Table 4**), features and UMIs, with the proportion of mitochondrial transcripts being higher in sample 1 and lower in sample 3 (**Fig. 2a**). As expected, there was a positive correlation between the number of detected genes and the number of UMIs, whereas a negative correlation was observed between the expression levels of mitochondrial genes and both gene counts and UMI counts, with no major differences between samples (**Fig. 2c**). Quality control filtering was performed by removing nuclei with mitochondrial transcripts representing more than 10% of the total counts, and upper and lower thresholds for UMI and number of unique genes were defined for each sample (see Methods). After filtering, initial clustering analyses revealed three clusters that were mostly composed of cells from a single sample that did not show expression of typical retinal cell marker genes (**Supplementary Table 2**). Since these nuclei most likely correspond to either low quality cells or non-retinal cells, they were removed from the dataset. The final object was composed of 17,438 cells containing 23,489 features (unique genes). There were no major differences in the number of unique genes, UMI count (number of transcripts) or mitochondrial feature levels between samples (**Fig. 2b, d**). However, sample 3 yielded a higher number of cells (**Table 4**), which can be attributed to minor technical differences in retinal dissection and/or library preparation.

**Table 4.** Final cell quantification and sequencing statistics. “Unfiltered nuclei” refers to the number of barcodes after importing the STAR files to Seurat, excluding cells with less than 200 UMIs. All other parameters refer to the final object.

| Sample | Sample 1 | Sample 2 | Sample 3 | Total |
| --- | --- | --- | --- | --- |
| Unfiltered nuclei | 69,723 | 52,932 | 64,138 | 186,793 |
| Filtered nuclei | 4,869 | 4,130 | 8,439 | 17,438 |
| Filtered features | 22,472 | 22,458 | 23,014 | 23,489 |
| Mean UMIs per cell | 4,527 | 3,716 | 3,486 | 3,831 |
| Median UMIs per cell | 3,333 | 2,234 | 2,359 | 2,572 |
| Mean features per cell | 2,153 | 1,885 | 1,852 | 1,944 |
| Median features per cell | 1,892 | 1,434 | 1,502 | 1604 |
| Mean percent of mitochondrial features | 3.55 | 2.66 | 2.13 | 2.65 |
| Median percent of mitochondrial features | 3.06 | 2.05 | 1.64 | 2.05 |

**Figure 2.**
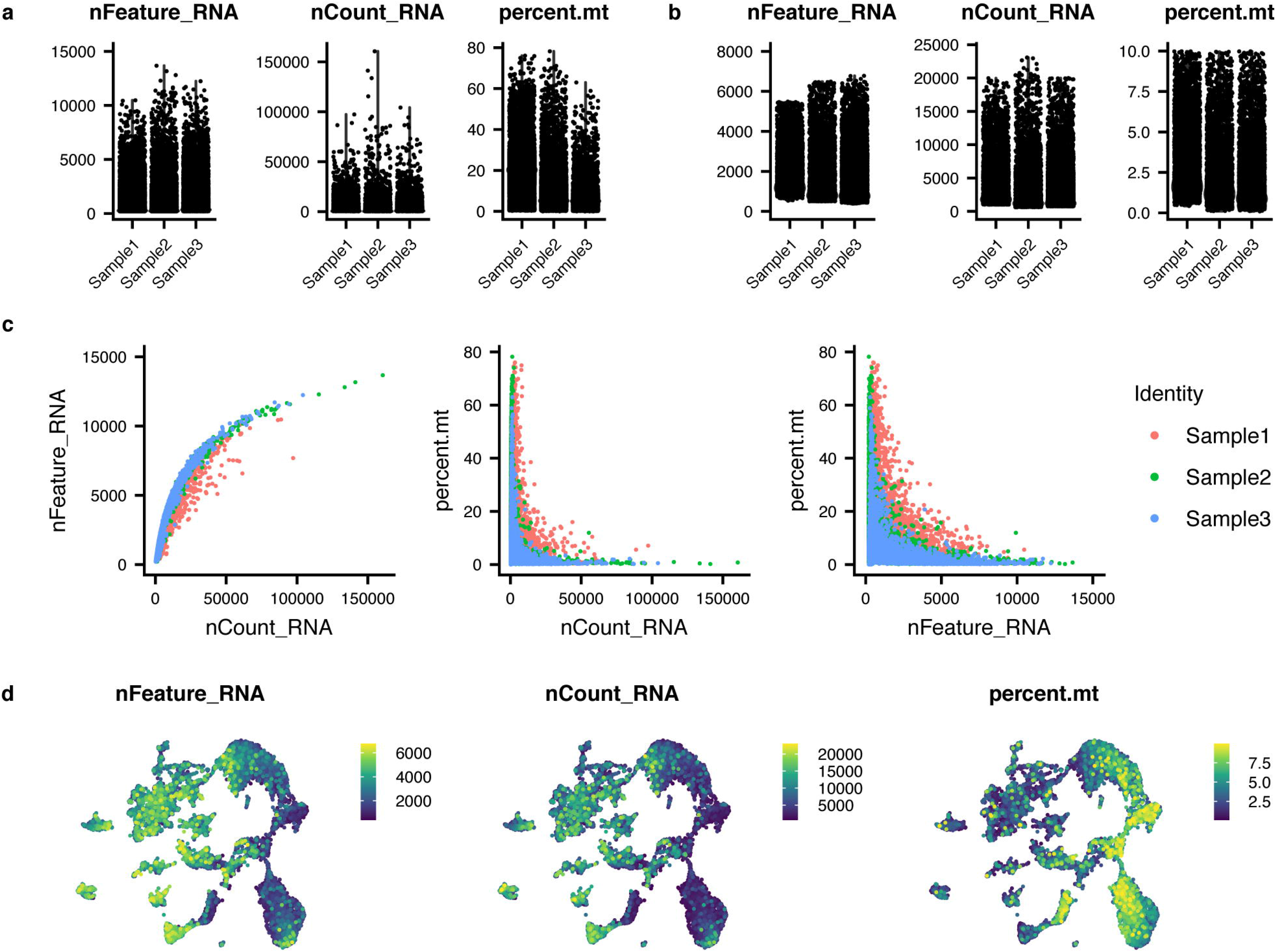
Quality control for snRNA-seq data. (**a, b**) Violin plots showing: left, the number of detected unique genes (“nFeature_RNA”); middle, the number of unique molecular identifiers (UMIs, “nCount_RNA”); and right, percentage of mitochondrial genes (“percent.mt”) in each cell from each sample before (**a**) and after (**b**) quality control. Note that all mitochondrial features were removed from the analyses after quality control (see Methods). (**c**) Scatter plots showing the relationship between gene counts and UMI counts (left), UMI counts and percentage of mitochondrial genes (middle), together with gene counts and percentage of mitochondrial genes (right) in the three samples before quality control. (**d**) Gene counts (left), UMI counts (middle) and percentage of mitochondrial genes (right) mapped onto the UMAP.

### Cluster annotation

Unsupervised clustering of filtered cells revealed 22 clusters (**Fig. 3a**), all of which were present in generally similar proportions in the three samples (**Fig. 3b**), with cells from the three retinas showing similar distributions across the UMAP (**Fig. 3c**), thus confirming the similarity of the samples. To annotate the clusters, we analysed the expression levels of known marker genes from other species (see Methods) for the major retinal cell classes or subclasses (**Fig. 4a, b**). This allowed us to divide cells into the six major retinal cell classes (PRs, HCs, BCs, ACs, RGCs and MG), as well as to identify two clusters of retinal progenitor cells (RPCs) and a single cluster of oligodendrocytes (OLs) (**Fig. 3a**).

**Figure 3.**
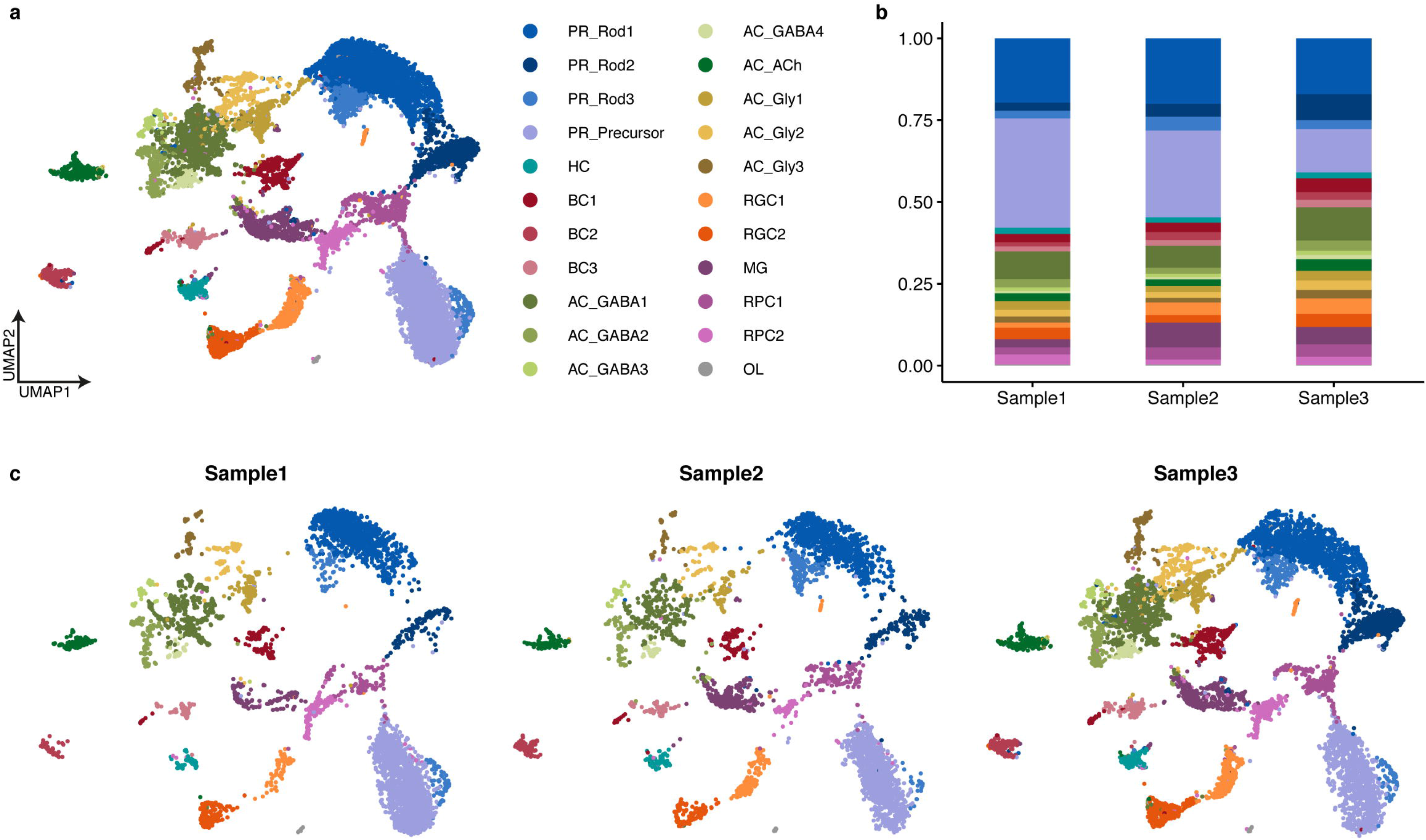
Clustering of snRNA-seq data reveals cell heterogeneity in the juvenile catshark retina. (**a**) UMAP of retinal cells showing their unbiased assignment to the 22 clusters identified in this study. For abbreviations, see **Table 3**. (**b**) Barplots showing the proportions of each cluster for each sample. (**c**) Separate UMAPs of cells from each sample, coloured according to their assigned identities.

**Figure 4.**
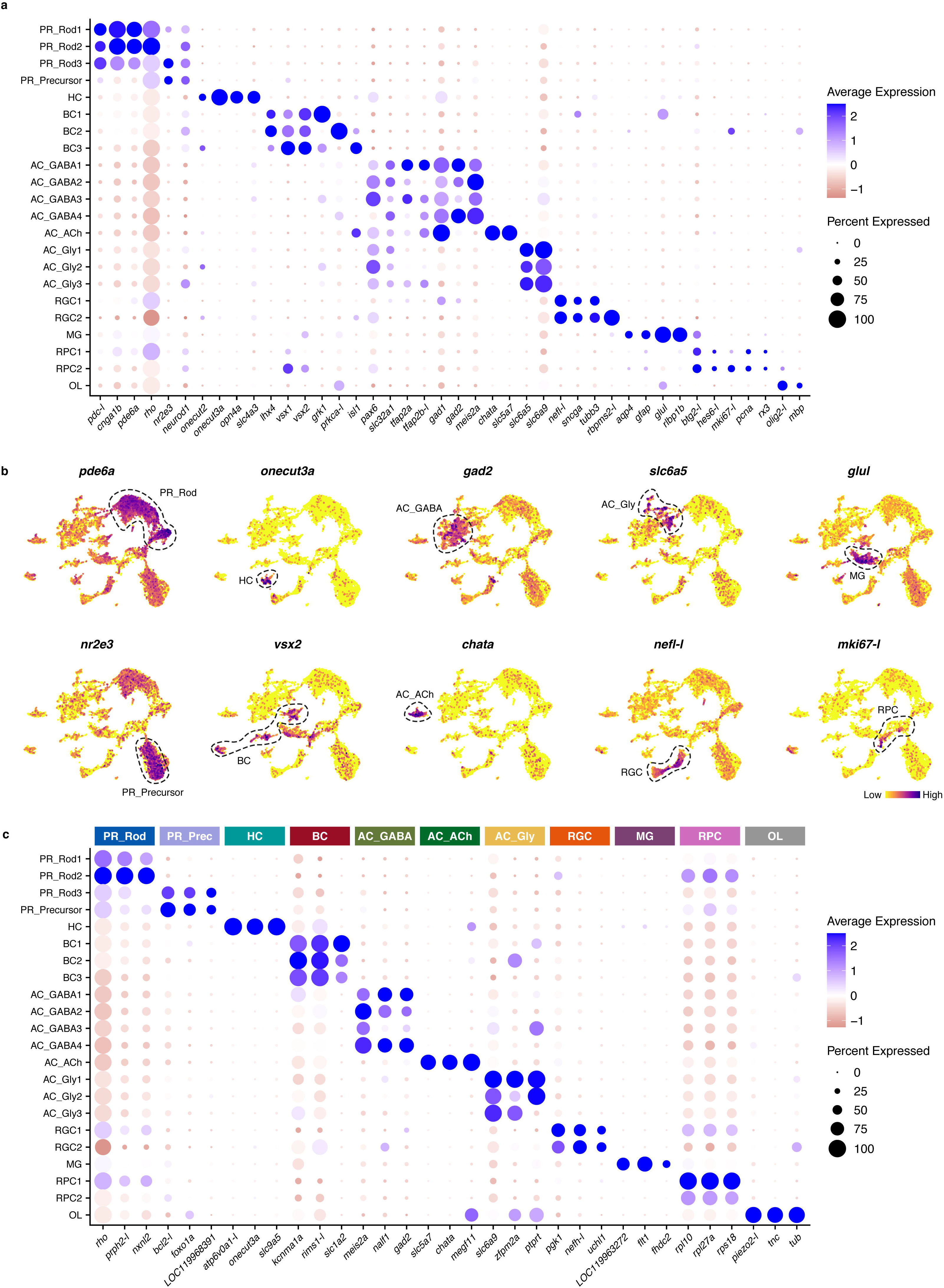
Expression of marker genes used for cluster annotation. (**a**) Dot plot showing expression levels of marker genes selected for identification of retinal cell classes (cluster annotation). (**b**) UMAPs showing expression of selected canonical marker genes. (**c**) Dot plot showing expression of the top three differentially expressed genes from class-level groupings (see text and **Table 3** for details). See Methods and **Supplementary Table 3** for information on gene nomenclature.

Three clusters could be identified as rod PRs by the expression of genes such as *rho, pdc-l* or *pde6a*^63^. Expression of established cone PR marker genes (not shown) was either present in rod clusters or virtually non-existing. This is in accordance with previous observations that *S. canicula* has either scarce cones^64^ or a pure-rod retina^46,65^ (reviewed in Ferreiro-Galve *et al*.^46^), similar to some other benthic sharks^65,66^ and the skate^67^. A fourth cluster that did not show high levels of most rod markers but showed significant expression of *nr2e3* and *neurod1*, known markers of rod precursors/progenitors^68^, was thus annotated as PR precursor cells. Since rod PRs usually comprise a single cell type^33,38,69^ (with some exceptions, such as those of the amphibian retina^70^), the four PR clusters identified here could correspond to different states of PR differentiation, but further research is needed to confirm this.

Regarding retinal interneurons, a single cluster of HCs (expressing the canonical marker *onecut3a*^33,71^) and 3 clusters of BCs (expressing *vsx1* and *vsx2*^11,14,72^, as well as high levels of *isl1* in one of the BC clusters, confirming previous immunohistochemical results^73^) could be identified in our dataset. We also found eight clusters of ACs, most of which showed high levels of *pax6*, confirming previous results^74^. The AC clusters could be further divided into three subclasses according to the expression of small molecule neurotransmitter-associated genes. Thus, we found four clusters of GABAergic ACs expressing *gad1* and/or *gad2*^18,75^; a single cluster of cholinergic (starburst) ACs, which show high levels of *chata* (similar to those described in *Squalus*^76^); and three clusters of glycinergic ACs, which show high levels of glycine transporters *slc6a5* and *slc6a9*^18^. Furthermore, two clusters of RGCs (expressing canonical markers, such as *nefl-l*^15^) could be recognised.

MG were represented by a single cluster with high levels of *rlbp1b*^11,77^ and *glul* (the gene encoding glutamine synthetase, whose expression in *S. canicula* MG has previously been described by means of immunohistochemistry^73,78^). Two clusters of RPCs could also be identified by the expression of cell proliferation markers (*mki67-l*^19,79^, *pcna*^45,46,78^) and typical markers of retinal progenitors (*btg2-l, rx3*^12^). Finally, a small cluster of OLs was identified by high levels of *olig2-l* and *mbp* expression^13,33^, probably corresponding to cells from the optic nerve head or the optic nerve fibres layer, where myelinated fibres are present in other elasmobranchs^42^. A remarkable absence from our dataset are microglial cells, which are known to be present in the innermost retinal layers of the postnatal retina of *S. canicula*^80^. This is most likely due to microglia being lost in the nuclei dissociation or quality control filtering processes.

To further validate cluster annotation and find novel markers specific to the *S. canicula* retinal cell classes, we identified marker genes (differential gene expression analyses vs the remaining cells in the dataset), both for individual clusters and for clusters grouped into classes/subclasses (**Table 3**). **Fig. 4c** shows the top marker genes for each class-level group, including both established and novel marker genes. For a complete list, see Data Records. Overall, the expression of marker genes confirms that our atlas comprises all major cell classes expected in the juvenile catshark retina, although some low abundant cell types (e.g. microglia) could be absent from the dataset. Thus, this dataset will provide a groundwork for studies on cell type diversity in the shark retina, allowing a better understanding of vertebrate retinal evolution and development.

## Usage Notes

All the code used to analyse the dataset is available in GitHub (see Code Availability). Raw .*fastq* files are standard Illumina sequencing files for 10x Genomics single-cell RNA sequencing libraries, and as such they can be processed using any typical single-cell analysis software (STAR^50^, as in our work, or Cell Ranger^81^, Kallisto^82^, Alevin^83^, etc.). STAR output files for each sample can be loaded into Seurat using the “ReadSTARsolo” function, and then merged into a single Seurat object using the “merge” function. Finally, the *Scyliorhinus_retina*.*rds* file is a Seurat object with the processed dataset that can be loaded into R with “ReadRDS”.

## Code Availability

The code used to process the raw sequencing files and generate all the results presented in this study can be found in https://github.com/Roslin-Aquaculture/SHARK_retina.

## Acknowledgements

This work was supported by grant ED431C 2021/18 funded by Xunta de Galicia to E.C., and grant PID2020-115121GB-I00 funded by MICIU/AEI/10.13039/501100011033 to A.B.-I. N.V.-V. was supported by grant FPU21/03076 funded by Ministerio de Ciencia, Innovación y Universidades. D.R. was supported by the Axencia Galega the Innovación (GAIN, Xunta de Galicia) as part of the Oportunius programme, and by BBSRC Institute Strategic Grants to the Roslin Institute (BBS/E/20002172, BBS/E/D/30002275, BBS/E/D/10002070 and BBS/E/RL/230002A). S.S. gratefully acknowledges an NSERC PDF award. This project has received funding from the European Research Council (ERC) under the European Union’s Horizon 2020 research and innovation programme (VerteBrain to H.K., grant agreement no. 101019268).

## Author contributions

Conceptualisation: N.V.-V., I.H.-N., A.B.-I., E.C. Data acquisition: I.H.-N., P.C.-P., S.S., P.R.V., F.H.-S., X.Y., C.S., J.S., S.M. (provided early access to the catshark genome), H.K. (supervision), D.R. Data analysis: N.V.-V., P.C.-P., S.S., P.R.V., F.L., D.R., E.C. Data interpretation: N.V.-V., P.C.-P., A.B.I., E.C. Writing (original draft): N.V.-V., P.C.-P., S.S., D.R., A.B.-I., E.C. Writing (review and editing): N.V.-V., I.H.-N., P.C.-P., S.S., P.R.V., S.M., H.K., F.A., D.R., A.B.-I., E.C. Funding acquisition: H.K., D.R., A.B.-I., E.C. Project supervision: E.C.

## Tables

**Supplementary Table 1**. Genes used in cell cycle scoring.

**Supplementary Table 2**. Differentially expressed genes from removed clusters.

**Supplementary Table 3**. Gene nomenclature.

